# Prophages divert *Staphylococcus aureus* defenses against host lipids

**DOI:** 10.1101/2022.01.27.478126

**Authors:** Biyang Zhou, Amit Pathania, Deepak Pant, David Halpern, Philippe Gaudu, Patrick Trieu-Cuot, Andressa Dias-Leao, Charlotte Pagot, Audrey Solgadi, Alexandra Gruss, Karine Gloux

## Abstract

Phages are ubiquitous in bacteria, including clinical *Staphylococcus aureus*, where Sfi 21/Sa3 phages often integrate into the *hlb* gene, encoding Hlb sphingomyelinase. The integration acts as a rapid regulatory switch of Hlb production. Our findings suggest that Sfi 21/Sa3 prophages and Hlb activity affect *S. aureus* fitness by modulating the incorporation of the toxic linoleic acid (C18:2) from serum into the bacterial membrane. This process relies on C18:2 derived from 1,3-diglyceride, facilitated by the FakB1 kinase subunit. Palmitic acid (C16), primarily released from serum through Hlb activity, competes for FakB1. This mechanism contributes to adaptation to AFN-1252, an antibiotic inhibiting the fatty acid synthesis pathway (anti-FASII). Since *S. aureus* relies on exogenous fatty acids for growth, AFN-1252 treatment leads to increased proportion of membrane C18:2. Moreover, Hlb inhibition, whether *via* prophage insertion, gene inactivation, or enzyme inhibition, delays *S. aureus* adaptation, resulting in higher proportionof C18:2 in the membrane. This study sheds light on the role of lipid environments in infections, and may contribute to the accurate prediction of infection risks and therapeutic efficacy. Furthermore, given that both anti-FASII and Hlb inhibitors enhance C18:2 incorporation, they represent potential agents for combined strategies against *S. aureus*.

## Introduction

Lipids are crucial for bacterial fitness and play a pivotal role in modulating interactions with the host (1–4). However, the impact of environmental lipids on bacterial pathogenesis is often overlooked. Lipids vary in structure from simple short hydrocarbon chains to complex molecules, including triglycerides (TGs), phospholipids (PLs), esterified sterols, and sphingolipids. Fatty acids (FAs), the fundamental constituents of complex lipids, differ in chain length, side chains, and the number and positions of double bonds. In humans, pathogens encounter diverse lipid environments influenced by genetics and diet, which can correlate to risk of infection (2,3,5,6). Bacterial lipases degrade host-derived lipids into free fatty acids (FAs), which a wide array of bacteria can scavenge and incorporate in their membranes (7–12). Notably, *Staphylococcus aureus*, a major human pathogen, incorporates exogenous FAs, such as those present in serum lipids (8,13). Once liberated, these FAs are incorporated using the FakAB system, which includes the FakA FA kinase and the binding proteins FakB1 and FakB2, which have preferences for saturated and unsaturated FAs, respectively (14–16). *S. aureus* lacks desaturase functions and the ability to synthesize unsaturated FA (17). However, the incorporation of monounsaturated FAs and polyunsaturated FAs (PUFAs) can significantly affect both bacterial membrane functions and interactions with the host (18–20).

Prophages of the Sfi 21/Sa3 family are the most prevalent phages integrated in staphylococcal genomes. They are present in over 90% of human clinical isolates (21–25). These prophages have an insertional hotspot in the *hlb* gene (*hlb*-conversion) encoding the *S. aureus* neutral sphingomyelinase C (Hlb, EC 3.1.4.12, also known as hemolysin B), which hydrolyzes sphingomyelins (Fig. 1A) (21,24). The significance of Sfi 21/Sa3 prophages for *S. aureus* adaptation to the human host is attributed to the immune evasion cluster (IEC) encoded by the prophage, and to the regulation of Hlb activity (26,27). Sfi 21/Sa3-prophages act as active lysogens, meaning they can excise from the *hlb* gene in response to environmental signals, such as biocides or reactive oxygen species, without killing bacteria (26,27). The released phage DNA (termed episome) can facilitate rapid adaptation through reinsertion in the same site. Although the roles of sphingomyelinases and IEC in virulence are well studied ((25,28), for review), the impacts of these prophages on *S. aureus* physiology are less understood. Furthermore, Sfi 21/Sa3-prophages are implicated in the host switch between animals and humans (29,30), making their physiological impact on bacteria important for a One Health approach against *S. aureus*.

**Figure 1.**
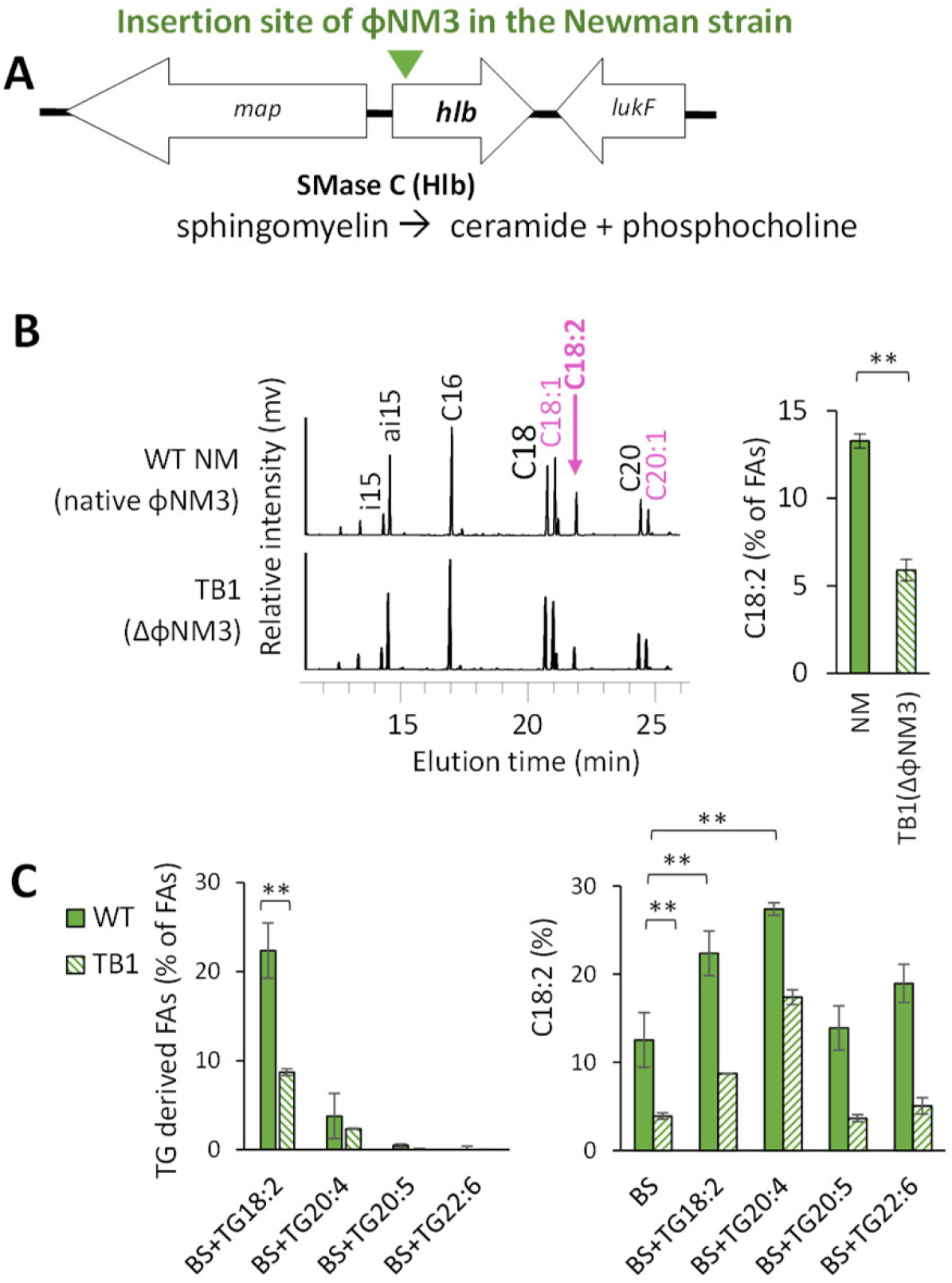
*hlb*-converting prophage enhances C18:2 incorporation from serum and triglycerides (TGs). (A) The insertion site of phage ɸNM3 in the *hlb* gene, which encodes the enzyme sphingomyelinase Hlb, results in a truncation at the enzyme N-terminus position 56 of the 330 amino acid sequence. The gene *map* encodes an extracellular adherence protein of broad specificity (Eap/Map); and the gene *lukF* encodes a protein from the leukocidin/hemolysin toxin family. (B) *hlb*-conversion increases the incorporation of linoleic acid (C18:2) from mouse serum. The wild type Newman strain (WT NM, containing the *hlb*-converting ɸNM3 prophage) and the TB1 strain (lacking ɸNM3 and with an intact *hlb* gene) (see Table 1) were grown for 2 hours in BHI medium supplemented with 10% mouse serum. FAs were extracted from PLs and analyzed by gas chromatography. Left, representative FA profiles are shown with y-axis representing the relative FA abundance (mV response) at indicated peak positions. Endogenously produced FAs are in black, while FAs not synthesized by *S. aureus* but incorporated from serum are in purple. Right, histograms depict the relative amounts of C18:2 incorporated into PLs. (C) *hlb*-conversion specifically affects the incorporation of C18:2 from TGs into PLs. The WT NM and TB1 strains were cultured in BHI medium supplemented with 10% bovine serum and enriched with 30µM of triarachidonin (TG20:4), trieicosapentaenoin (C20:5) or tridocosahexaenoin (C22:6). Data are presented as mean +/- standard deviation from independent experiments (n=4). Statistical significance was determined using the Mann Whitney test on incorporated FAs. **, p≤0.01.

**Table 1.**
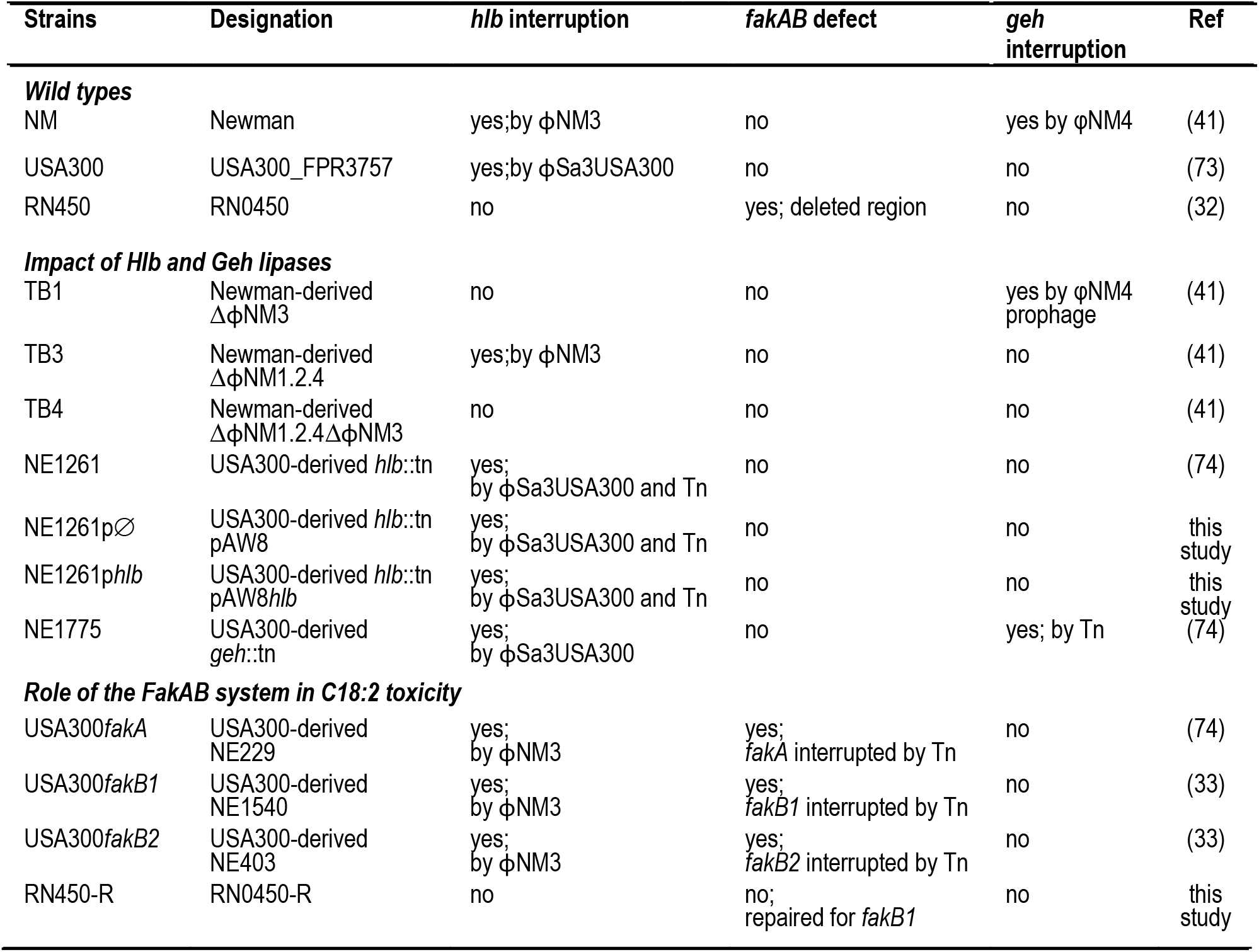
Strains and phages.

In this study, we observed that interruption of the *hlb* gene by the φNM3 prophage (from the Sfi 21/Sa3 family) results in increased incorporation of the bactericidal PUFA linoleic acid (C18:2) into the membrane PLs of *S. aureus*. We investigated the mechanism by which the presence of an *hlb*-converting prophage affects the incorporation of C18:2, which is primarily released from TGs (31). Our findings demonstrate that a TG hydrolysis intermediate leads to high incorporation of C18:2 *via* the FakB1 protein subunit. This unexpected binding to FakB1 results in competition with the C16 FA released from sphingomyelins for incorporation into PLs. Furthermore, we showed that inhibition of FASII and/or Hlb activities exacerbates the incorporation and toxicity of C18:2 in PLs, reducing *S. aureus* fitness.

## Materials and methods

### Bacterial strain

*S. aureus* USA300 FPR3757 (methicillin-resistant, referred to as USA300), and NCTC 8325 derivative strains RN450 and RN450-R, were used in this study (Table 1). RN450, also designated as 8325-4, is a phage-free derivative of the NCTC 8325 strain (32) and has a deficient *fakB1* gene due to a natural deletion. RN450-R is an RN450 derivative repaired for *fakB1* as described below. Isogenic Newman strains differentiated by the presence of prophages, wild type (referred to as WT NM), TB3 (Δɸ11), TB1 (ΔɸNM3) and TB4 (Δɸ11,ΔɸNM3), were provided by the Schneewind laboratory (Department of Microbiology, University of Chicago). The WT NM has 4 prophages, including fNM3 from the Sfi 21/Sa3 subfamily inserted in *hlb*, and fNM4 inserted in the *geh* encoding a TG lipase (21). The TB3 strain has only the *hlb*-converting prophage fNM3. The Nebraska USA300 library of transposon insertions (University of Nebraska Medical Center) was generously supplied by BEI resources (33). The Nebraska library derivatives used in this work contained insertions, verified by PCR, in the following genes: SAUSA300_0320 (*geh*), SAUSA300_1973 (*hlb*), SAUSA300_1119 (*fakA*), SAUSA300_0733 (*fakB1*), SAUSA300_1318 (*fakB2*). The different strains used in this study, their *hlb* and *geh* status (intact or interrupted genes), and prophage insertions are presented in Table 1.

### Construction of an hlb complemented strain

A PCR fragment containing the *hlb* promoter and the open reading frame SACOL_RS10470 (−355 from the ATG start codon and +40 after the TAG stop codon) from the *S*. *aureus* subsp. *aureus* COL, was cloned between EcoRI and SmaI sites of plasmid pAW8 (34). The primers pairs used for the PCR were 5’-TTGCCGGAATTCTGCAACTTAATTATAGCCAGACTTTC-3’ and 5’-CATCAACCCGGGCGTCCTTTTAGAACGAAGCAAG-3’. Genomic coordinates of the 1388 bp cloned segment were 2063371-2064758. This plasmid, named p*hlb*, was used for complementation experiments.

### Construction of the RN450-R strain, repaired for fakB1

The *S. aureus* RN450 *fakB1* gene lacks a 483-bp internal segment. This deletion occurs in the entire NCTC 8325 lineage, removing 56% of the 867-bp functional gene (Fig. S1). To repair this deletion, a 1,939-bp DNA fragment containing a functional *fakB1* was amplified to repair RN450, as described for other NCTC 8325 derivatives (35).

### Growth media and conditions

Brain heart infusion (BHI) was the base medium for *S. aureus* cultures. Bacteria were cultured aerobically at 37°C with a starting OD_600_ of 0.02, starting from 2 to 5 hour precultures prepared in BHI. Mouse or adult bovine sera (Clinisciences, France) were added at 10% final concentration. Stocks of trilinolein (TG18:2, 50 mM), triarachidonine (TG20:4, 10mM), trieicosapentaenoin (TG20:5, 50mM), tridocosahexaenoin (TG22:6, 50mM), 1,2-dilinoleoylglycerol (1,2DG, 50 mM), 1,3-dilinoleoylglycerol (1,3DG, 50 mM), palmitic acid (C16, 100 mM), (Larodan Fine Chemicals, Sweden), orlistat (38 mM, MedChemExpress, France), and the anti-FASII antibiotic AFN-1252 (1 mg/ml, MedChemExpress, France) were prepared in DMSO.

In the absence of FASII inhibitor, FA profiles and C18:2 incorporation were determined from bacteria recovered after short growth times (OD_600_ between 1 and 4) to avoid total consumption of C18:2. Bacterial growth was consequently presented as variation of OD_600_ per hour (OD_600_ /h). To follow *S. aureus* adaptation to an anti-FASII, bacteria were pre-cultured for 4 hours in BHI medium supplemented with 10% mouse serum (Eurobio, France), and inoculated at OD_600_ 0.02 in the same medium containing 0.5 µg/ml AFN-1252. Bacteria were cultured for 17 hours at 37°C in 96-well microtiter plates using a Spark spectrophotometer (Tecan), and OD_600_ was measured every 10 minutes. Growth results are presented as bacterial growth kinetics as a function of time.

### Hlb substrate, inducer and inhibitor

Sphingomyelin was from chicken egg yolk (Sigma-Aldrich; St. Louis, MO) and prepared as 50 mg/ml stock solution in chloroform/methanol (1:1). Hlb sphingomyelinase belongs to the neutral sphingomyelinase family, which requires a lipid-dependent activation by phosphatidylserine. Phosphatidylserine (60 mM, bovine brain, Na salt, Larodan Sweden) was supplied in chloroform. GW4869 (Selleckchem.com), a specific inhibitor of the Hlb conserved domain involved in this lipid activation (36,37), was prepared at 1.7 mM in DMSO.

### Determination of S. aureus fatty acid profiles

FAs profiles were performed as described previously (9). Briefly, culture aliquots were taken during exponential growth at low OD_600_ (between 1 and 4). After sample preparation, extraction and trans-esterification of FAs (9), GC separation was performed by injection of the FA methyl esters in a split-splitless mode on an AutoSystem XL gas chromatograph (Perkin-Elmer) equipped with a ZB-Wax capillary column (30 m x 0.25 mm x 0.25 mm; Phenomenex, France) and a flame ionization detector. FAs were identified based on their retention times and coinjection with purified methyl esters FA standards (Mixture ME100, Larodan, Sweden). Data were recorded and analyzed using a TotalChrom Workstation (Perkin-Elmer). FA peaks were detected between 12 and 40 minutes of elution. Results are shown as representative gas chromatograph profiles, or as FA peak areas expressed as percentage of the total areas of detected peaks.

### Detection and characterization of diglycerides, ceramides, and sphingomyelins

Lipid extractions were performed as described (38) with modifications (39). Briefly, freeze-dried supernatants (10ml) of culture (OD_600_ from 3 to 4) were extracted with 9.5 ml of chloroform-methanol 0.3% NaCl (1:2:0.8 v/v/v) at 80°C for 15 min and vortexed for 1 h at room temperature (RT). After centrifugation at 4000 rpm for 15 min (RT) supernatants were collected and debris were re-extracted with 9.5 ml of the same mixture, and vortexed for 30 min. After centrifugation, supernatants were pooled and 2.5 ml each, of chloroform and 0.3% NaCl solutions, were added and mixed. Phase separation was achieved by centrifugation at 4000 rpm for 15 min (RT). The upper phase was discarded and the collected chloroform phase was evaporated to dryness under a nitrogen stream and stored at −20°C. Lipids were identified following the previously described method (40). They were separated by using a normal phase HPLC (U3000 ThermoFisher Scientific) equipped with an Inertsil Si 5µm column (150 x 2.1 mm I.D.) from GL Sciences Inc (Tokyo, Japan). Lipids were identified by mass-spectrometry negative ionization and MS^2^/MS^3^ fragmentations (LTQ-Orbitrap Velos Pro). Lipid spectra were analyzed using Xcalibur^TM^ software (ThermoFisher Scientific, version 4.2.47).

### Statistics analysis

Graphs were prepared using Microsoft Excel software. Means and standard deviations are presented for culture growth (OD_600_ or OD_600_/h), FA percentages, and percentages of FA elongated *via* FASII. Statistical significance was determined by unpaired, nonparametric Mann Whitney tests, as recommended for small sample sizes or by non-parametric paired test Wilcoxon Signed Rank test for dependent samples.

## Results

### The hlb-converting prophage ɸNM3 increases C18:2 incorporation from serum into S. aureus phospholipids

Two isogenic Newman strains of *S. aureus*, WT and TB1, which differ by the presence of the *hlb*-converting prophage ɸNM3 (Table 1)(41), were grown in control BHI medium. FAs present in their PLs were extracted and analyzed. Both strains displayed identical FA profiles corresponding to endogenous FAs produced by FASII activity (Fig. S2), indicating that the ɸNM3 prophage does not affect *S. aureus* FAs present in membrane PLs in this culture condition. However, supplementation with 10% mouse serum led to the incorporation of exogenous PUFAs, which *S. aureus* cannot synthesize (17). Notably, only the proportion of C18:2 in PLs was significantly affected by the presence of ɸNM3, being two-fold higher in the WT than in TB1 (Fig. 1B). To confirm this observation, we tested adult bovine serum, which contains more diverse PUFAs, including C18:2, α-linolenic acid (C18:3 ω-3), dihomo-γ-linolenic acid (20:3 ω−6), and arachidonic acid (C20:4 ω-6) (Fig. S3). Since diverse PUFAs are found in the human body, mostly transported as TGs, we enriched cultures with different TGs, each carrying a single distinct type of long-chain PUFA, as frequently found *in vivo*: C18:2, C20:4 ω-6, eicosapentaenoic acid (C20:5 ω-3), or docosahexaenoic acid (C22:6 ω-3). FA profiles from WT and TB1 strains grown with these diverse sources demonstrated that only two PUFAs were incorporated into *S. aureus* PLs: C18:2 and, to a very low extent, C20:4 (Fig. 1C). Furthermore, C18:2 incorporation was the most affected by the presence of ɸNM3 prophage (Fig. 1 B & C). Enrichment with 30µM trilinolein (TG18:2 in Fig. 1C), a TG source containing three C18:2 molecules, further increased the proportion of C18:2 in PLs in both strains, indicating that the prophage maintains its effects despite the enrichment. Interestingly, enrichment with triarachidonin (TG20:4), a TG source of C20:4, also increased C18:2 incorporation in both strains, even more than enrichment with TG18:2 (Fig. 1C). Although the mechanism remains unclear, these results suggest a prophage strategy specifically modulating C18:2 incorporation in the *S. aureus* membrane. This effect is unexpected since serum sphingomyelins are poor sources of C18:2 (18,42). To verify this, we analyzed sphingomyelins present in the BHI-mouse serum medium used in our experiments. We therefore questioned how ɸNM3 is connected to C18:2 incorporation into *S.aureus* PLs.

### C18:2 from 1,3-dilinoleoylglycerol is highly incorporated and toxic in S. aureus

*S. aureus* triglyceride lipases can produce both 1,2 and 1,3 diacylglyceride intermediates from TG hydrolysis, as illustrated by the TG18:2 example (Fig. 2A). We analyzed the impact of 1,2-dilinoleoylglycerol (1,2DG) and 1,3-dilinoleoylglycerol (1,3DG) on *S. aureus* growth and their efficiency as substrates for C18:2 incorporation into PLs (Fig. 2B). Since 1,3 DG is preferentially released by Geh lipase activity (43,44)(Fig. 2A), the experiment was conducted using the USA300 *geh^+^* and *geh^−^*strains (Table 1). We observed that 1,3DG was highly incorporated in *S. aureus* PLs compared to 1,2DG (Fig. 2B). Additionally, 1,3DG was significantly more toxic in the *geh*^+^ strain than in the isogenic *geh^−^*strain (Fig. 2B). However, the resulting proportion of C18:2 and elongated forms into PLs is minimally affected, suggesting that it is not the determining factor of toxicity. However, both the hydrolysis position on the glycerol and the *geh* status are crucial for toxicity of the released C18:2. As the culture condition of the first experiment showed *hlb*-conversion effect on C18:2 incorporation, we further determined if 1,3-diglycerides can be released by *S. aureus* in the presence of serum and if they can be, as 1,3DG, sources of PUFAs (Fig. 1B). We used a USA300 *fakA* mutant to block FA incorporation (14) and increase the likelihood of finding 1,3-diglyceride intermediates. Mass-spectrometry negative ionization and MS^2^/MS^3^ fragmentations identified 1,3-diglycerides in the supernatant after growth of the *fakA* mutant in the presence of 10% mouse serum (Fig. 2C). Orlistat, an inhibitor of Geh activity (45), reduced the levels of both 1,3-diglycerides and the 1,3-diglycerides/1,2-diglycerides ratio in the supernatant (Fig. 2C). Characterization of the 1,3-diglycerides species demonstrated that orlistat also decreased relative abundance of 1,3-diglycerides with polyunsaturation (Fig. 2D). In conclusion, in the presence of serum, 1,3-diglycerides intermediates are released by the bacterial lipase and serve as sources of PUFAs. Since more than 85% of *S. aureus* isolates are *geh*^+^ (1), most *S. aureus* strains may release 1,3-diglyceride sources of PUFAs.

**Figure 2.**
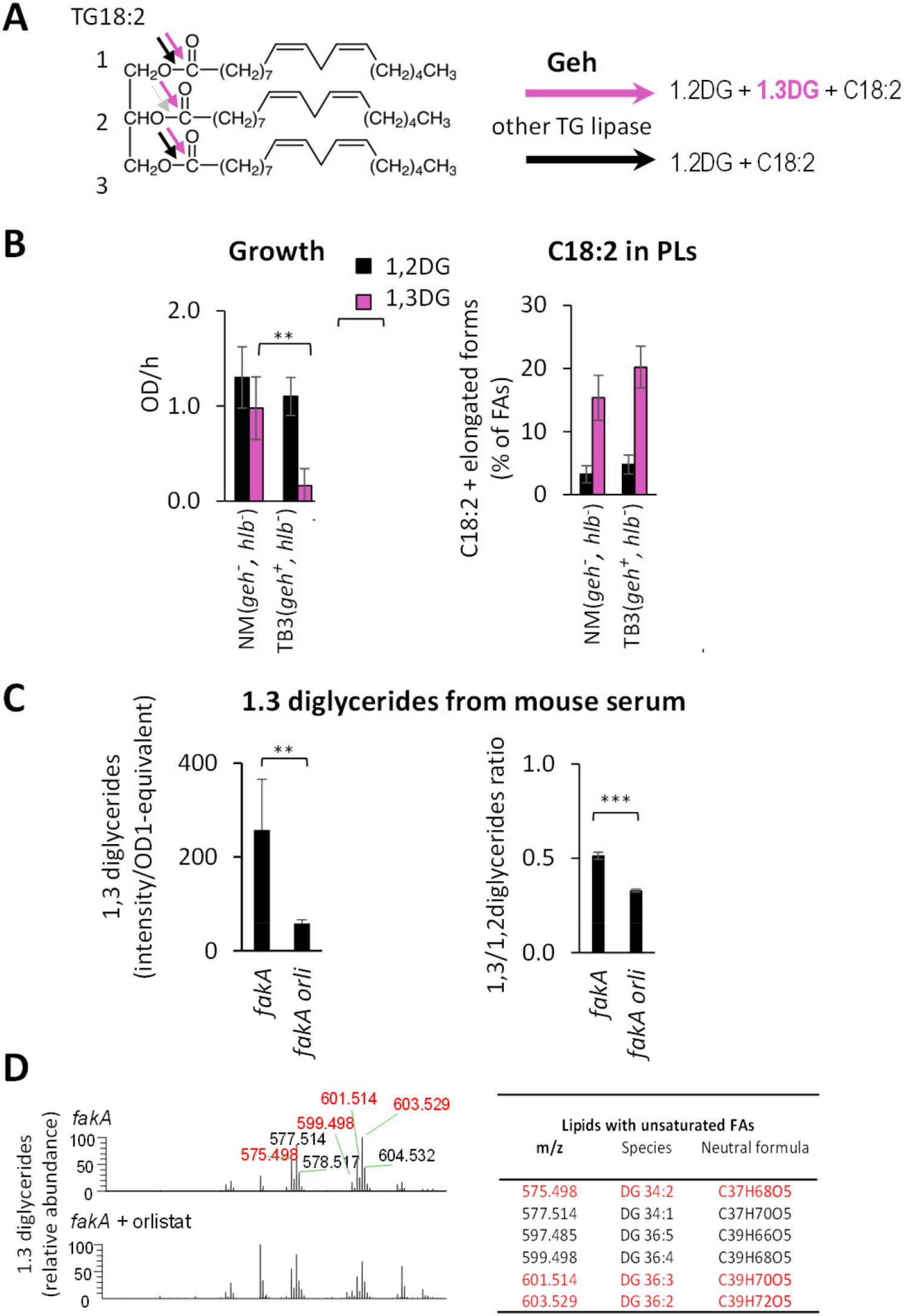
The role of 1,3-diglycerides in C18:2 toxicity on *S. aureus*. (A). The non-selective cleavage of TGs by *S. aureus* Geh activity results in both 1,2- and 1,3-diglycerides (1,2DG and 1,3DG from TG18:2). Most lipases preferentially cleave at position 1, releasing 1,2-diglycerides, as illustrated for 1,2DG from TG18:2 In contrast, Geh can cleave at both positions, thus releasing 1,3-diglycerides (42, 43). (B) The 1,3DG supplementation results in high incorporation of C18:2 into PLs. WT NM (*geh*::tn) and TB3 (intact *geh*) strains (see Table 1) were cultured for 2 hours in BHI medium supplemented with 10 µM of either 1,2DG or 1,3DG. The relative amounts of incorporated C18:2 (C18:2 and its elongated forms) were determined as in Figure 1B and are presented as FAs derived from the diglycerides. Notably, C18:2 was more efficiently incorporated and toxic when derived from 1,3DG compared to 1,2DG, with TB3 (intact *geh*) showing particular sensitivity. (C) *S. aureus* lipase generates 1,3-diglycerides. The USA300 *fakB1*::tn mutant was cultured in BHI medium with 10% mouse serum, with or without the addition of 30µM orlistat. Lipids present in the supernatant were extracted and analyzed using normal phase HPLC. Diglycerides were identified by their retention times and through comparison with pure diglyceride standards. (D) The 1,3-diglycerides comprise polyunsaturations. The 1,3-diglyceride species were identified by mass-spectrometry using negative ionization and MS^2^/MS^3^ fragmentations. Data are presented as mean +/- standard deviation from independent experiments (n=4). Statistical significance was determined by Mann Whitney test on incorporated C18:2 or by the non-parametric Wilcoxon Signed Rank test on 1,3 diglycerides intensities per OD_600nm_=1 equivalent and the 1,3-/1,2-diglyceride ratio.**, p≤0.01. ***, p≤0.001.

### hlb protects S. aureus from 1,3DG-derived C18:2, and promotes C16 incorporation over C18:2

Although sphingomyelins are not sources of C18:2 in our experiments, we investigated the potential impact of the *hlb* gene on C18:2 incorporation into PLs. For this purpose, we used the NE1261 strain, a USA300 FPR3757 with an inactive *hlb* gene due to both conversion and transposon insertion, and performed *hlb* complementation. A plasmid carrying an intact *hlb* (p*hlb*), from the *S. aureus* COL strain, was constructed and introduced into the NE1261 strain (Table 1). NE1261p*hlb* and the control strain (empty plasmid, NE1261pØ) were grown in BHI supplemented with 10% mouse serum and 10 µM 1,3DG, without or with sphingomyelin (36)(see Material and Methods). In the absence of sphingomyelin, *hlb* complementation had little or no effect on growth rate; remarkably however, C18:2 incorporation into PLs was significantly inhibited (Fig. 3A). Moreover, sphingomyelin supplementation led to significantly improved growth and concomitant poor incorporation of C18:2 from 1,3DG into PLs. Surprisingly, *hlb* complementation also significantly inhibited the elongation of both saturated and unsaturated FAs (Fig. 3B). As C18:2 was not transferred into PLs, it may be maintained in free form intracellularly, and inhibit the enoyl reductase enzyme (FabI) in FASII (31). Since sphingomyelins are potential FA sources (Fig. 3. B), we hypothesized that FAs can be released and compete with C18:2 for incorporation into PLs. An intermediate step involving a ceramidase would be necessary to release such competitive FAs. Although *S. aureus* is not known to produce ceramidase, it can be present in serum (46). To confirm this possibility, we searched for ceramide intermediates in the BHI-mouse serum used for cultures and in the supernatant after growth of *S. aureus* USA300. Palmitic acid (C16) is a common sphingomyelin constituent in human blood (42,47). We identified C16 as the main FA in sphingomyelins under BHI-mouse serum conditions of our first experiment, and ceramide intermediates with C16 acyl sources were found in culture supernatants (Fig. 3B). Furthermore, *hlb*-complementation increased C16 levels in PLs (Fig. 3C). Based on these findings, the effect of the *hlb*-converting prophage on membrane FAs likely reflects an interference between C16 and C18:2 incorporation. We further investigated the underlying mechanism.

**Figure 3.**
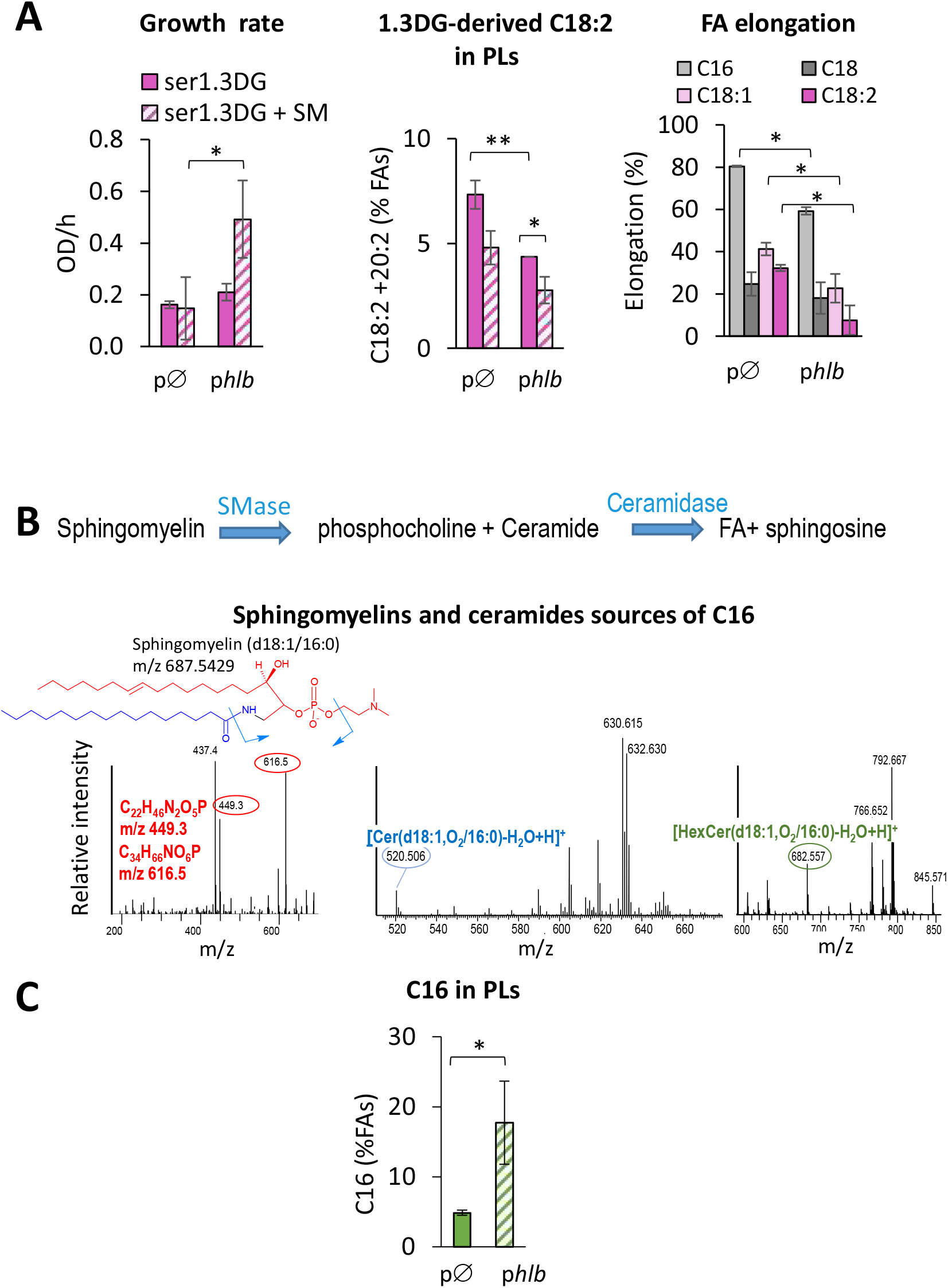
Plasmid-driven expression of *hlb* affects *S. aureus* fitness, inhibits FASII, and increases C16 in PLs. (A) *hlb*-complementation protects from 1,3DG-derived C18:2 incorporation and toxicity but inhibits FA elongation. The NE1261 mutant (Tn insertion in *hlb*) bearing an empty plasmid (pØ) or a plasmid carrying an intact *hlb* (p*hlb*) (Table 1) was grown for 3 hours in BHI medium supplemented with 10% mouse serum and 10 µM 1,3DG. Activated sphingomyelin (SM, 200 µM) was added as an Hlb substrate (see Materials and Methods). Left, growth is presented as OD_600_ increase per hour. Middle, incorporation in PLs of C18:2 and its elongated form C20:2. Relative amounts of incorporated FAs were determined as in Fig. 2B and expressed in percent of total FAs. Right, FA elongation by FASII determined from relative amounts of elongated forms from C16, C18, C18:1 and C18:2, and expressed as a percentage of total (FA + elongated) for each. (B) Sphingomyelins and ceramides from *S. aureus* culture supernatant are sources of C16. The USA300 strain was cultured in BHI medium in the presence of 10% mouse serum. Lipids were extracted, and sphingomyelins and ceramides were identified and characterized by NPLC-MS in positive atmospheric pressure chemical ionisation, with species confirmed by MS/MS. Left: The main sphingomyelin species deduced from NPLC and subsequent fragmentations was SM(d18:1/16:0). Detection threshold of SM species were around 0.025 mg/mL. Note that no sources of C18:2 were detected (only traces of SM(d18:1/18:1) and SM(d18:1/18:0)). Middle and Right: Ceramide intermediates containing C16 were detected in culture supernatants: a dihydroceramide (middle, Cer(d18:1,O2/16:0)), and a hexylceramides (right, HexCer(d18:1,O2/16:0)). (C) Plasmid-driven expression of *hlb* increases C16 incorporation from sphingomyelin. Cultured of NE1261 pØ; and p*hlb* strains and FAs analysis were performed as in (A). Total C16 incorporated in PLs was determined as for C18:2 in Fig. 1B and expressed as percent of total FAs. Data presented in histograms are means +/- standard deviations from independent experiments (n=3). Statistical significance was determined by the Mann-Whitney test on OD/ h, incorporated FAs or elongation percentages. **, p≤0.01, *p≤0.05.

### Incorporation of 1,3DG-derived C18:2 requires the kinase subunit FakB1 and is inhibited by C16

The substrate selectivity of the Fak kinase complex is determined by the FakB1 and FakB2 FA binding proteins, which reportedly bind to saturated and unsaturated FAs, respectively (14–16). We confirmed these phenotypes using the USA300 *fakA*, *fakB1*, and *fakB2* mutants (Table 1) grown in media including saturated and unsaturated free FAs (Fig. S4A&B). When challenged with TG18:2, the *fakA* mutant did not incorporate C18:2 confirming that phosphorylation is required for incorporation of C18:2 from TG18:2 into PLs (Fig. S5A). Interestingly, both *fakB1* and *fakB2* mutants failed to show selectivity; both mutants displayed similar capacities to incorporate C18:2 from TG18:2 (Fig. S5A, left). This indicates that while the incorporation of free C18:2 is FakB2-dependent, incorporation from TG18:2 can utilize both FakB proteins.

We further investigated whether the diglycerides released from TG18:2 by *S. aureus* lipases share this FakB binding flexibility. Compared to the TG18:2, incorporation of 1,2DG-derived C18:2 was significantly reduced in the *fakB2* mutant (Fig. S5A right) aligning with the reported FakB2 binding preference for free unsaturated FAs (14–16). In contrast, incorporation of 1,3DG-derived C18:2 was higher in the *fakB2* mutant than in the *fakB1* mutant (Fig. 4). Moreover, growth of the *fakB1* mutant was strongly impaired compared to the wild-type strain (Fig. 4). These results demonstrate that *fakB1* mediates the incorporation of 1,3DG-derived C18:2 and likely provides protection against the high toxicity of this diglyceride. This unexpected role of *fakB1* suggests a competition with the Hlb-derived C16, a known FakB1 ligand (14). To test this hypothesis, we added C16 in the presence of TG18:2 or 1,3DG. In both the USA300 WT and the *fakB1* mutant strains, the addition of free C16 led to a slight decrease in C18:2 incorporation from TG18:2 or 1,3DG (Fig. 4 and Fig. S5B). In stark contrast, in the *fakB2* mutant (which only has FakB1 available), C16 addition almost completely halted incorporation of C18:2 from both sources. Since 1,3DG is an intermediate product of TG18:2 hydrolysis, exogenous C16 may compete with 1,3DG-derived C18:2 for binding to FakB1. Interestingly, the protective effect of FakB1 against the 1,3DG was not observed in the presence of C16 (Fig. 4 middle). The presence of C16 may lead to free C18:2, which is toxic to bacteria and inhibits FASII activity, in accordance with the observed effect of *hlb* on FA elongation (Fig.3A right). This may represent a type of toxicity that is distinct from toxicity due to C18:2 incorporation in PLs. To confirm the direct role of *fakB1* in the incorporation of 1,3DG-derived C18:2, we used the RN450 strain, which carries an in-frame deletion in *fakB1* (Fig. S1). This deletion affects two FA binding sites and the phosphotransfer reaction site, resulting in a non-functional protein. This strain is cured of prophages and carries an intact *hlb* gene (Table 1). We constructed a derivative strain, designed RN450-R, in which *fakB1* is repaired (Table 1). As expected from the above results, 1,3DG-derived C18:2 was more efficiently incorporated by RN450-R than by the *fakB1* defective RN450 strain (Fig. 5). In RN450, the presence of C16 had no effect on the incorporation of 1,3DG-derived C18:2, likely due to the absence of FakB1. In contrast, C16 strongly inhibited the incorporation of 1,3DG-derived C18:2 in the RN450-R strain (Fig. 5), confirming a direct role of *fakB1* in mediating the competition between 1,3DG-derived C18:2 and C16 incorporation into PLs. C16 supplementation did not significantly affect growth of RN450-R, suggesting strain-specificities regarding the competition impact on the toxicity of the released C18:2. However, these observations collectively identify *fakB1* as encoding the preferred binding protein for 1,3DG-derived C18:2 and as the key mediator of the competitive incorporation of C18:2 and C16.

**Figure 4.**
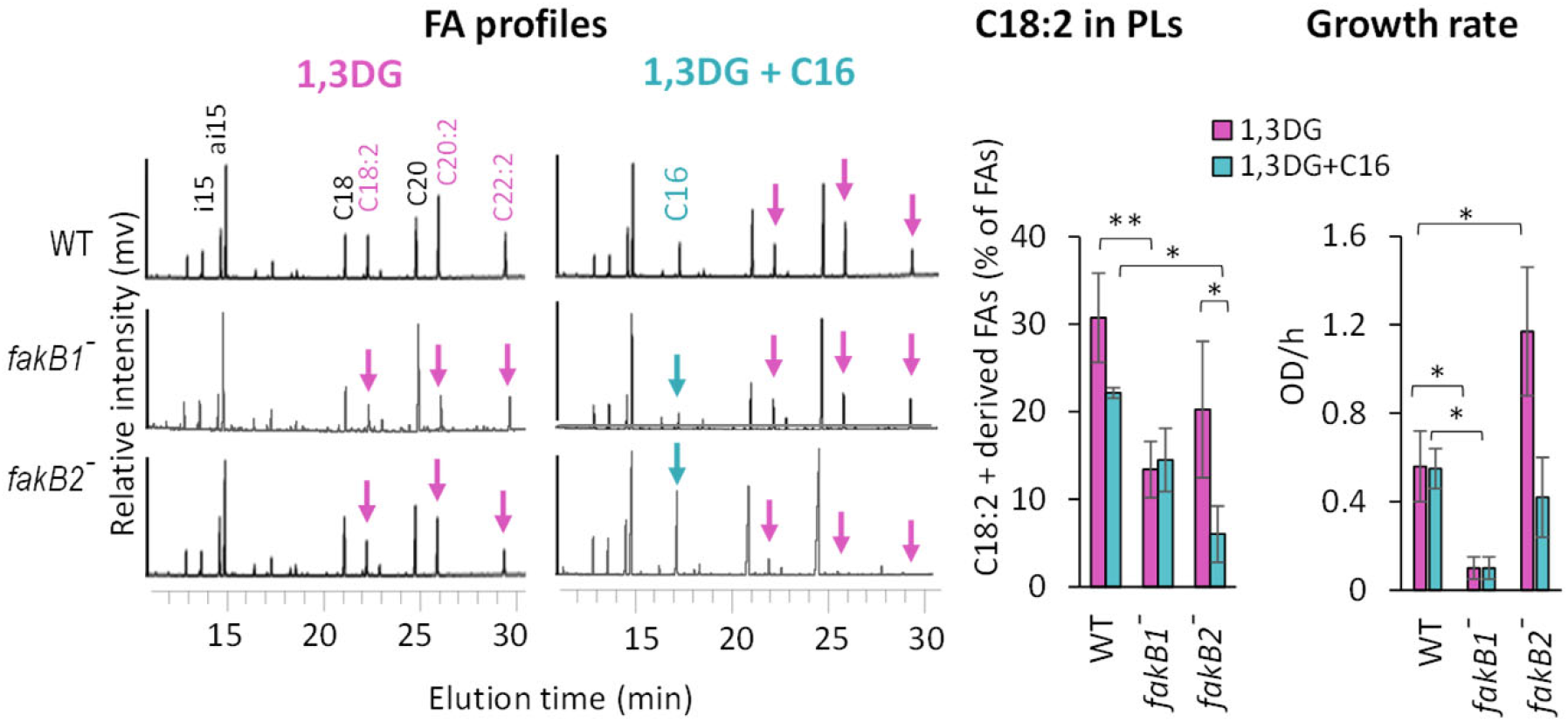
FakB1 mediates the incorporation of 1,3DG-derived C18:2, competing with C16 incorporation. USA300 (WT), *fakB1::tn*, and *fakB2::tn* mutant strains (see Table 1) were cultured for 2 hours in BHI medium supplemented with 10 µM 1,3DG, with or without 20 µM C16. FA profiles were analyzed as described in Figure 1B. Bacterial growth and incorporated C18:2 were measured as in Figure 3A. In the FA profiles, purple arrows indicate C18:2 and its elongated forms (C20:2 and C22:2). Growth is presented as OD_600_ per hour (OD/h). Data in histograms are means +/- standard deviations from independent experiments (n=5). Statistical significance was determined by the Mann-Whitney test on C18:2 incorporation from 1,3DG and OD_600_/h.**, p≤0.01; *, p≤0.05.

**Figure 5.**
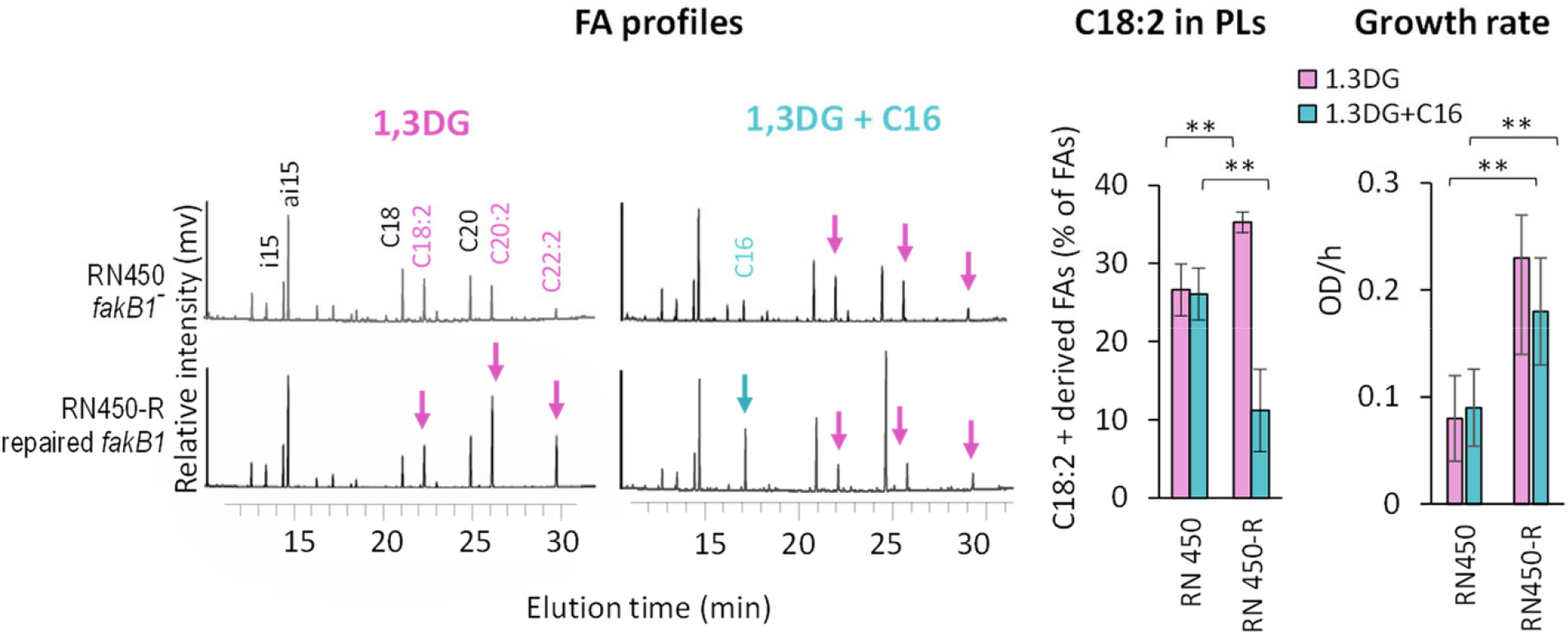
Restoration of *fakB1* enhances incorporation of 1,3DG-derived C18:2 and competition with C16. RN450 (*fakB1*^−^) and RN450-R (*fakB1*^+^) strains (see Table 1) were cultured for 3 hours in BHI medium supplemented with 10 µM 1,3DG, with or without 40 µM C16. FA profiles were determined as described in Figure 1B. Bacterial growth and incorporated C18:2 were measured as described in Figure 3A. In the FA profiles, purple arrows indicate C18:2 and elongated forms (C20:2 and C22:2). Growth is presented as OD_600_ per hour (OD_600_/h). Data in histograms are means +/- standard deviations from independent experiments (n=5). Statistical significance was determined by the Mann-Whitney test on C18:2 incorporation from 1,3DG and OD_600_/h.**, p≤0.01.

### Inhibitors of FASII and Hlb increase C18:2 incorporation and limit S. aureus adaptation

In the presence of an antibiotic targeting FASII (anti-FASII), *S. aureus* adapts by incorporating exogenous FAs, including unsaturated FAs (8). Consequently, bacterial growth becomes exclusively reliant on exogenous FAs under this condition, likely increasing levels and toxicity of PUFA in the membrane. We therefore analyzed the potential synergistic effects of disabling both FASII and Hlb on *S. aureus* growth. AFN-1252 was the FASII inhibitor; this drug targets FASII enoyl reductase enzyme FabI and is effective and safe against skin infections (48,49). GW4869 inhibits *S. aureus* sphingomyelinase activation (36,37). Growth of TB3 and TB4 isogenic strains (Table 1) was compared in BHI-mouse serum medium supplemented with 30µM 1,3DG, with or without GW4869. When FASII was active, TB3 (*hlb*-converted) exhibited slightly decreased growth compared to TB4 (intact *hlb*), and GW4869 reversed the growth advantage of TB4 (Fig. 6A left). Addition of the FASII inhibitor AFN-1252 significantly slowed growth. As reported, *S. aureus* adapted to AFN-1252 in the presence of serum (8) but the TB4 strain displayed a markedly shortened latency period prior to adaptation compared to TB3 (outgrowth <2h instead of 12h) (Fig. 6A right). Furthermore, the addition of GW4869 did not significantly change the growth of TB3 but delayed that of TB4. These results demonstrate that Hlb activity limits inhibitory effects of AFN1252. We determined FA profiles from these cultures and analyzed the C18:2 percentage in PLs. AFN-1252 increased the incorporation of C18:2 into the PLs of the TB4 strain, especially in the presence of GW4869. The C18:2 percentage in membrane PLs of TB4 was increased fourfold by AFN-1252 treatment (Fig. 6B). These results suggest that Hlb activity accelerates adaptation to anti-FASII by limiting C18:2 incorporation and toxicity in membrane. Safety in humans was demonstrated for both anti-FASII antibiotic AFN1252 and the Hlb inhibitor GW4869 (48,50), suggesting that they may be good candidates for combinatory strategies against *S. aureus*.

**Figure 6.**
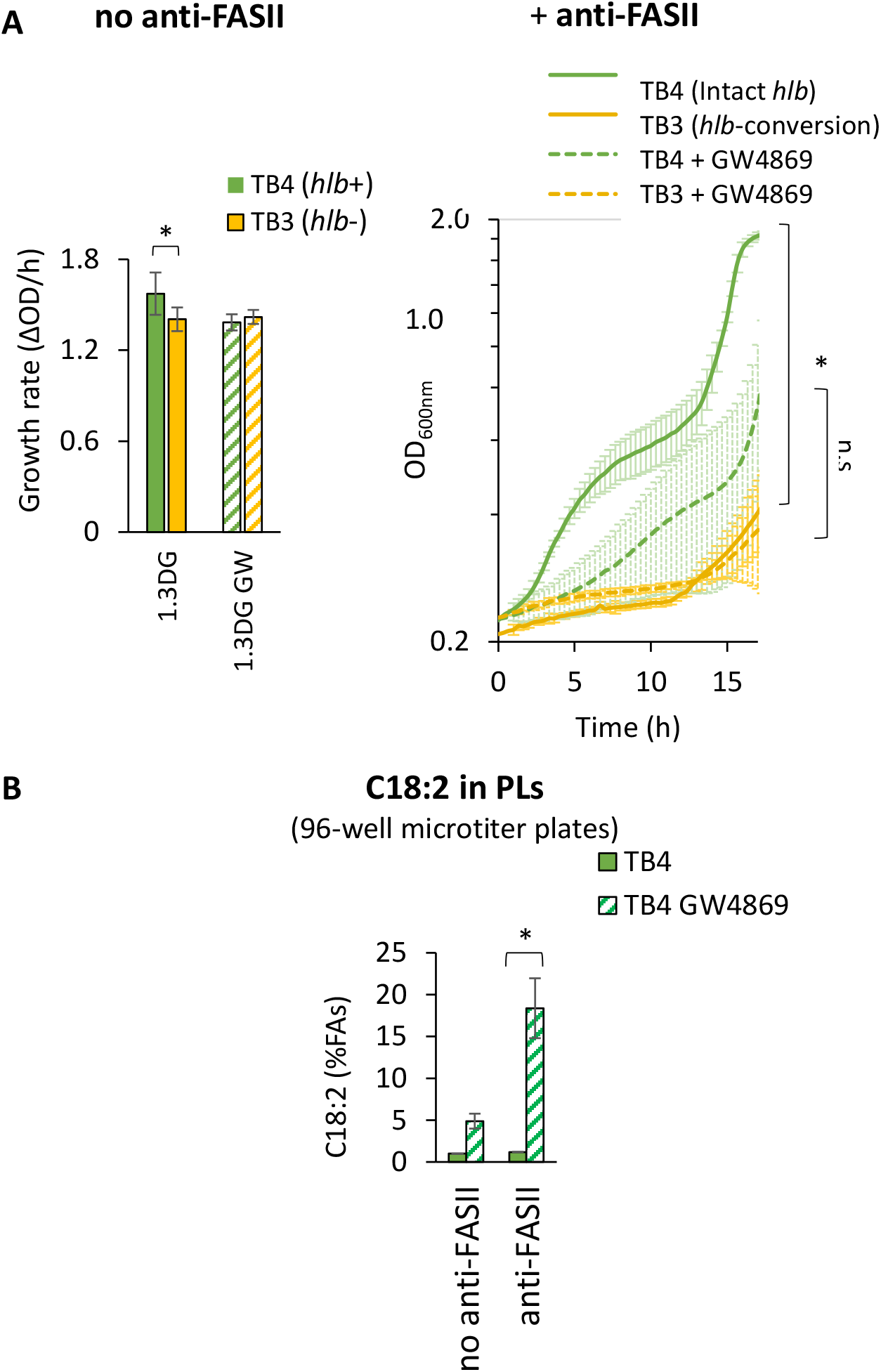
Hlb activity and anti-FASII treatments enhance C18:2 incorporation in *S. aureus* PLs. TB3 (фNM3 in *hlb*) and TB4 (no phage) strains (see Table 1) were cultured in BHI medium supplemented with 10% mouse serum, 30 µM 1,3DG, and with or without 50µM GW4869 (an inhibitor of Hlb activity). Upper left: bacterial growth rates in the absence of anti-FASII treatment, performed in glass tubes. Upper right, growth kinetics in the presence of 0.5µg/ml AFN-1252 (a FASII inhibitor), performed in 96-well microtiter plates, with OD600 measured every 10 minutes. Bottom: C18:2 incorporation with or without AFN-1252. C18:2 incorporation was determined in the TB4 and TB3 strains as described in Figure 2B. Data are means +/- standard deviations from independent experiments (n=4). Statistical significance was determined by the Mann-Whitney test on OD_600_/h and by the Wilcoxon Signed Rank test on incorporated C18:2; *, p≤0.05. n.s: not significant.

## Discussion

This study reveals the crucial role of *hlb*-converting prophages and Hlb sphingomyelinase activity in the lipid metabolism of *S. aureus* and its adaptation to the presence of an anti-FASII agent (Fig. 7A & 7B). We first showed that Hlb activity unexpectedly inhibits C18:2 incorporation into the bacterial membrane. The connection was demonstrated to be the FakB1 kinase subunit: FAs released from TGs and those released from sphingomyelins both compete for this function (Fig. 7A). This mechanism involves a TG-derived 1,3-diglyceride, which leads to abundant C18:2 incorporation *via* FakB1 binding, in competition with sphingomyelin-derived C16. FakB1 affinity for C18:2, derived specifically from 1,3DG, and not from free C18:2, nor from 1,2-diglyceride-derived C18:2, remains to be understood. The Hlb/C18:2/FakB1connection may represent a natural Achilles’ heel exploited by Sfi 21/Sa3 prophages to modulate bacterial fitness in response to PUFA toxicity.

**Figure 7.**
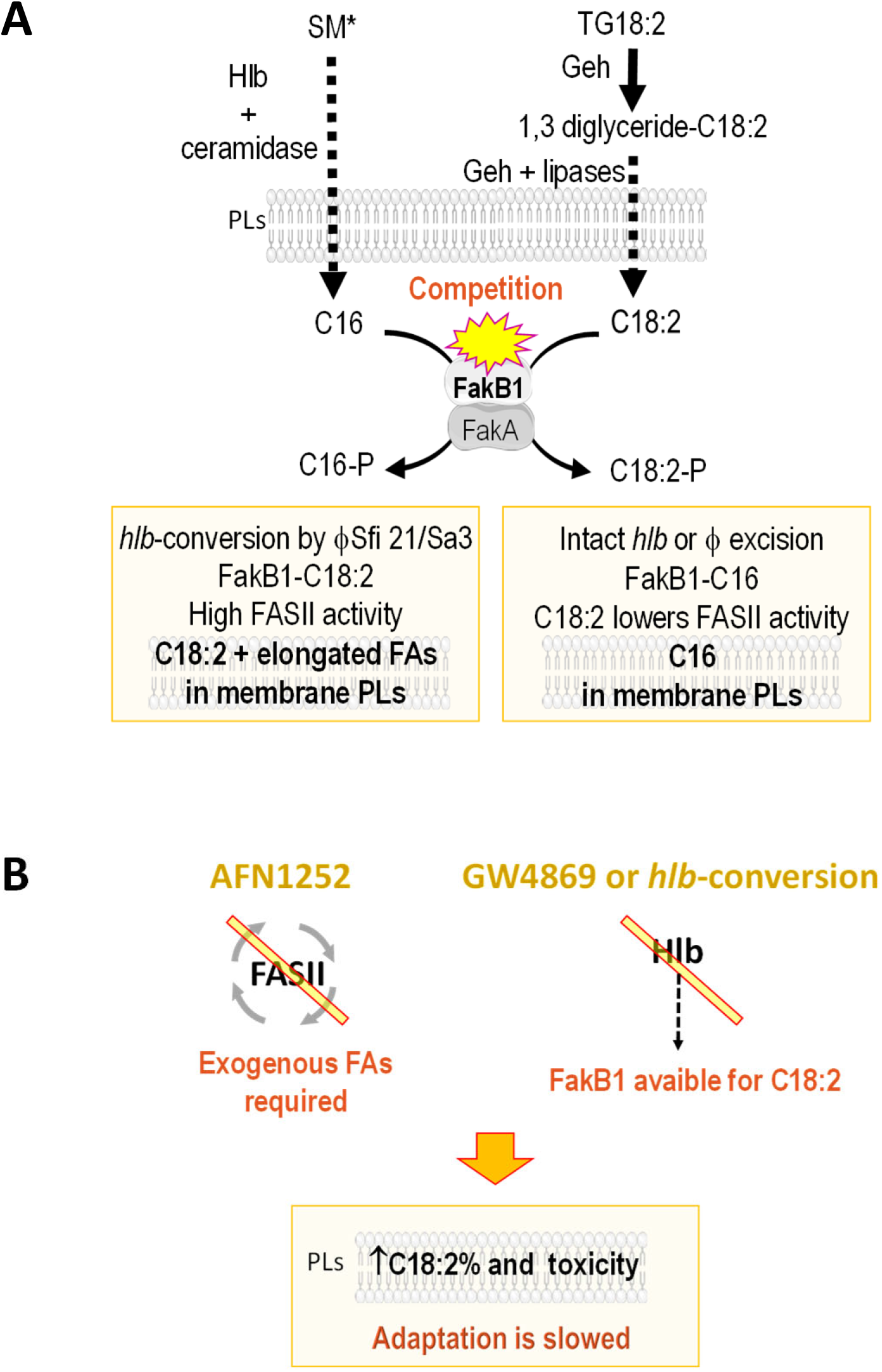
Mechanistic model of C18:2 toxicity on *S. aureus* and enhancement by FASII or sphingomyelinase inhibition. (A) Modulation of C18:2 incorporation by the *hlb*-converting prophage is based on 1,3-diglyceride and free FA competition for FakB1. The model highlights three major features: 1) The 1,3-diglyceride intermediate, released by *S. aureus* Geh lipase, leads to high incorporation and toxicity of C18:2 in the *S. aureus* membrane. 2) C18:2 derived from the 1,3-diglyceride binds to FakB1 for incorporation into PLs, in contrast to free C18:2 or that derived from 1,2DG, which is non-competitive. FakB1 is one of the two acyl-binding subunits of the FakAB kinase complex, which phosphorylate FAs and promotes membrane incorporation. 3) Hlb activity contributes to C16 release from serum sphingomyelins (SMs). C16 then binds to FakB1, preventing C18:2 binding and incorporation into PLs. The unbound C18:2 inhibits FASII (31), thereby hindering FA elongation and bacterial growth. Various stressors, as H_2_O_2_ or biocides, induce prophage excision into extrachromosomal episomes, allowing rapid phage excision and reinsertion, and control of Hlb activity (23,26). According to environmental Hlb and Geh substrates and the prophage insertion/excision status, *S. aureus* can alter its phospholipid composition and adapt to environmental C18:2. (B) Blocking FASII or sphingomyelinase activity exacerbates C18:2 incorporation, leading to reduced bacterial adaptation. Under anti-FASII conditions, S *aureus* relies on exogenous FAs for growth (8), incorporating PUFAs like C18:2, which are toxic when integrated into membrane PLs. Furthermore, inhibiting Hlb activity reduces competition for FakB1 binding, allowing greater C18:2 incorporation into PLs (see Fig. 7A). Therefore, inhibitors of both FASII and Hlb are potentially effective combinatory drugs to enhance the natural defense mechanism involving C18:2. SM*: activated sphingomyelin (see material and methods), TG-C18:2: triglyceride containing C18:2, 1,3 diglyceride-C18:2: 1,3-diglyceride containing C18:2.

TGs are main constituents in human fat and blood, and serve as major circulating reservoirs of C18:2 (51). During infection, both Geh and Hlb lipases are active (27,52). We speculate that C18:2/FakB1 binding and its competition with sphingomyelin-derived C16 may occur *in vivo*. Sfi21/Sa3 are *hlb-* converting prophages that act as Hlb regulatory switches in most clinical isolates (21–24,27,53). As active lysogens, these prophages may reversibly control this Hlb/C18:2/Fakb1 connection. This mechanism provides novel insights into the prevalence of *hlb*-converting prophages in clinical isolates and the role of the lipid environment in *S. aureus* adaptation to infection. Incorporation of PUFAs, such as C18:2, into PLs alters crucial functions that interact with the infecting host, including biofilm formation, secretion of virulence factors, and interference with immune defense (8,19,20,54–56). Furthermore, C18:2 and sphingolipid metabolism are key pathways affected by *S. aureus* to decrease macrophage efficacy (57). Therefore, control of the Hlb/C18:2/FakB1 connection by *hlb*-converting prophages may play an important role in *S. aureus* escape from innate immunity.

The results demonstrate that levels and balance of TGs and sphingomyelins, along with FA composition and position in TGs, are important factors for *S. aureus* fitness. TGs and sphingomyelins vary according to the state of infection (58,59), the infected site, and between individuals, based on diet and genetic makeup (51,58–60). Interestingly, several diseases associated with high risk of infection or severe evolution, as in Crohn’s disease, type 2 diabetes, obesity or cystic fibrosis, show dysbiosis in TGs and/or sphingomyelins (42,51,57,61–67). Novel strategies are particularly required against sepsis (68,69). Lipid emulsions are approved or under investigation for the intravenous nutrition of patients with sepsis ((https://clinicaltrials.gov/study/NCT03405870)(70)). The mechanism connecting TGs and SMs offers new hypotheses and possibly future lipid formulations against fatal outcomes.

The present work further shows that *hlb*-conversion limits adaptation to anti-FASII treatment (mechanistic model in Fig. 7A). Consistent with the aforementioned Hlb/C18:2/FakB1 connection, the absence of Hlb activity increases the incorporation of toxic C18:2 and delays anti-FASII adaptation. What can be inferred from previous *in vivo* studies of anti-FASII adaptation? *S. aureus* was demonstrated to efficiently adapt to anti-FASII in a mouse model (8). Since Hlb activity occurs during infection (for review (25)), we propose that prophage excision contributes to anti-FASII adaptation *in vivo*. Membrane fatty acid composition contributes to modulating the efficacy of other antibiotics such as daptomycin and vancomycin (71,72), making this Hlb/C18:2/FakB1 connection relevant for future drug screenings and development. Moreover, FASII and Hlb inhibitors promote greater C18:2 proportion in membrane PLs. Therefore, combining an Hlb inhibitor or anti-FASII upon PUFA enrichment may a have synergistic effect against *S. aureus*. Furthermore, a strategy incorporating a natural lipid-based mechanism can be promising in reducing drugs dosages and *S. aureus* survival. Genetic variation of prophages is a primarily driver in bacterial transmissibility between species (30). Considering the Hlb/C18:2/FakB1 connection, the TG/SM status should be an important determinant of interspecies propagation. In conclusion, the mechanism used by these prophages to divert *S. aureus* lipid metabolism is of paramount importance in addressing prevention and healthcare challenges.

## Data availability

This article contains supplemental data. All data supporting the findings of this study are available within this article and its supplemental data.

## Supplemental data

This article contains supplemental data.

## Acknowledgment

We gratefully acknowledge funding from the following French granting agencies: French Agence Nationale de la Recherche (StaphEscape project ANR-16-CE15-0013), the Fondation pour la Recherche Medicale (DBF20161136769), and the INRAE’s department MICA (Project LipStaph). We thank Myriam Gominet (Institut Pasteur) for the construction of the *S. aureus* RN450 *fakB1*-repaired strain. We are grateful to members of the MicrobAdapt team for stimulating discussion of this work, and to Charlotte Pagot for proofreading the manuscript.

## CRediT author statement

K. G., B. Z., A. P., D. P., C.P., A. DL., P. G., and D.H. investigation; K. G., A. G., A. P., P. G., and P. T-C. writing-review & editing; K.G., A. G., A. P., A. S. and P. T-C. methodology; K. G. conceptualization, validation, formal analysis, writing original draft, visualization, supervision, project administration. A.G. funding acquisition and resources.

## Founding sources

This work was supported by French granting agencies: French Agence Nationale de la Recherche (StaphEscape project ANR-16-CE15-0013), and the Fondation pour la Recherche Medicale (DBF20161136769).

## Abbreviations

FA: fatty acid
FakAB: kinase module including the FakA FA kinase, and FakB binding proteins
IEC: immune evasion cluster
Hlb or SMase: hemolysin B or sphingomyelinase C
orli: orlistat
PL: phospholipid
PUFA: polyunsaturated fatty acid
SM: sphingomyelin
TG: triglyceride
TG18:2: trilinolein
TG20:4: triarachidonine
TG20:5: trieicosapentaenoin
TG22:6: tridocosahexaenoin
1,2DG: 1,2-dilinoleoylglycerol
1,3DG: 1,3-dilinoleoylglycerol.

## Supplemental data

**Supplementary Figure S1.** FakB1 defect in the RN450 strain. Alignment of FakB1 sequences from Newman and NCTC 8325 strains. The NCTC 8325 lineage, comprising the RN450 strain, carries an in-frame deletion in *fakB1* (483-bp within the 867-bp intact *fakB1*). The non-deleted region of NCTC 8325 FakB1 shares 100% identity with the synonymous region in the Newman strain. Blue arrows indicate amino acids required for FA binding and for FakA-mediated phosphotransfer (1).

**Supplementary Figure S2.** FA profiles of *S. aureus* cultured in BHI medium are not affected by the prophage status. Four isogenic Newman strains, lysogenized or not by prophages as shown, were cultured in BHI. FAs were extracted and analyzed as described in Fig. 1B. Representative FA profiles and means of relative amounts (in purple) from three independent experiments are shown. Statistical significance was determined by the Kruskal-Wallis test to compare the FA compositions between the four strains. Profiles were not significantly different, p value> 0.05.

**Supplementary Figure S3.** FA profile from adult bovine serum. A representative profile determined as in Figure 1B is shown. This serum contains different PUFAs: linoleic acid (C18:2), α-linolenic acid (C18:3 ω-3), dihomo-γ-linolenic acid (20:3 ω−6), and arachidonic acid (C20:4 ω-6).

**Supplementary Figure S4.** Phenotype validation of *fakA, fakB1* and *fakB2* mutants. The USA300 wild type and mutant strains (Table 1) were cultured in BHI medium in the presence of exogenous FAs. Total membrane FAs were extracted, analysed and presented as in Fig. 1B. (A) FAs profiles from the WT strain and the *fak* mutant cultured in the presence of an FA cocktail (free C14:0, C16:0, and C18:1, 0.17 mM each). The three exogenous FAs and elongated forms are indicated in blue. Orange arrow indicates C18:1 elongation into C20:1. Blue crosses indicate that *fakA* is essential for FA incorporation and that *fakB2* is essential for C18:1 incorporation. Purple arrows indicate that *fakB1* is involved in incorporation of C14 and C16. (B) *fakB2* is required for incorporation of free C18:2. The WT strain and the *fakB2*::tn mutant were cultured in in the presence 10µM free C18:2. In profiles, C18:2 and its elongated forms are represented in blue. Note that free C18:2 is poorly incorporated in the *fakB2* mutant compared to the WT. Data presented are means +/- standard deviations from independent experiments (n=3). Statistical significance was determined by the Mann-Whitney test. *, p≤0.05.

**Supplementary Figure S5.** Roles of *fak* genes in the incorporation of C18:2 from TG18:2 or 1,2DG. Experiments and FAs profiles were performed as described in Figure 4, except that the USA300 WT and *fak* mutant strains (Table 1) were cultured in the presence of 30µM TG18:2 or 1,2DG, as sources of C18:2. (A) Representative FA profiles from *fakA, fakB1* and *fakB2* mutants cultured in the presence of TG18:2 or 1,2DG. Left: The *fakA* defective strain fails to incorporate C18:2 from TG18:2, while both *fakB1* and *fakB2* mediate incorporation of TG18:2-derived C18:2. Right: The *fakB2* gene, and not *fakB1*, affects C18:2 incorporation from 1,2 dilinolein (1,2DG). (B) C16 inhibits incorporation of C18:2 from TG18:2 *via* competition for FakB1. The *fakB2* mutant (encoding only *fakB1*) fails to incorporate TG18:2-derived C18:2 in the presence of C16. Growth rates presented as OD_600_ per hour suggest that FakB1 protects from C18:2 toxicity and that C16 inhibits growth when only FakB1 is active by releasing free C18:2 in *S. aureus*. TG18:2-derived C18:2 and elongated forms are indicated in blue, and blue crosses indicate their absence. Purple arrows indicate a C16 increase due to exogenous supply. Blue arrows indicate inhibition of TG18:2-derived C18:2 and elongated forms. Profiles are representative of independent experiments (n=3).Histograms are means +/- standard deviations from these experiments. Statistical significance was determined by the Mann-Whitney test.; *, p≤0.05.

