## supplementaries Figures for "Prophages divert *Staphylococcus aureus* defenses against host lipids"

```
                                                    carboxyl binding site
               10        20        30        40        50        60        70
           ....|....|....|....|....|....|....|....|....|....|....|....|....|....|
NCTC 8325  MKIAVMTDSTSYLSQDLIDKYNIQIAPLSVTFEDGK----------------------------------
Newman     MKIAVMTDSTSYLSQDLIDKYNIQIAPLSVTFDDGKNFTESNEIAIEEFYNKMASSQTIPTTSQPAIGEW

           carboxyl binding site
               80        90       100       110       120       130       140
           ....|....|....|....|....|....|....|....|....|....|....|....|....|....|
           ----------------------------------------------------------------------
           ITKYEMLRDQGYTDIIVICLSSGISGSYQSSYQAGEMVEGVNVHAFDSKLAAMIEGCYVLRAIEMVEEGY

               Phosphotransfer reaction
              150       160       170       180       190       200       210
           ....|....|....|....|....|....|....|....|....|....|....|....|....|....|
           ---------------------------------------------------------IIPEEKVRTKKRA
           EPQQIIDDLTNMREHTGAYLIVDDLKNLQKSGRITGAQAWVGTLLKMKPVLKFEDGKIIPEEKVRTKKRA

              220       230       240       250       260       270       280
           ....|....|....|....|....|....|....|....|....|....|....|....|....|....|
           IQTLEKKVLDIVKDFEEVTLFVINGDHFEDGQALYKKLQDDCPSAYQVAYSEFGPVVAAHLGSGGLGLGY
           IQTLEKKVLDIVKDFEEVTLFVINGDHFEDGQALYKKLQDDCPSAYQVAYSEFGPVVAAHLGSGGLGLGY

           ....|...
           VGRKIRLT
           VGRKIRLT
```

**Supplementary Figure S1.** FakB1 defect in the RN450 strain. Alignment of FakB1 sequences from Newman and NCTC 8325 strains. The NCTC 8325 lineage, comprising the RN450 strain, carries an in-frame deletion in *fakB1* (483-bp within the 867-bp intact *fakB1*). The non-deleted region of NCTC 8325 FakB1 shares 100% identity with the synonymous region in the Newman strain. Blue arrows indicate amino acids required for FA binding and for FakA-mediated phosphotransfer (1).


1. Cuypers M.G., Subramanian C., Gullett J.M., Frank M.W., White S.W., Rock C.O. (2019) Acyl-chain selectivity and physiological roles of *Staphylococcus aureus* fatty acid-binding proteins. *J Biol Chem.* **294**(1):38-49

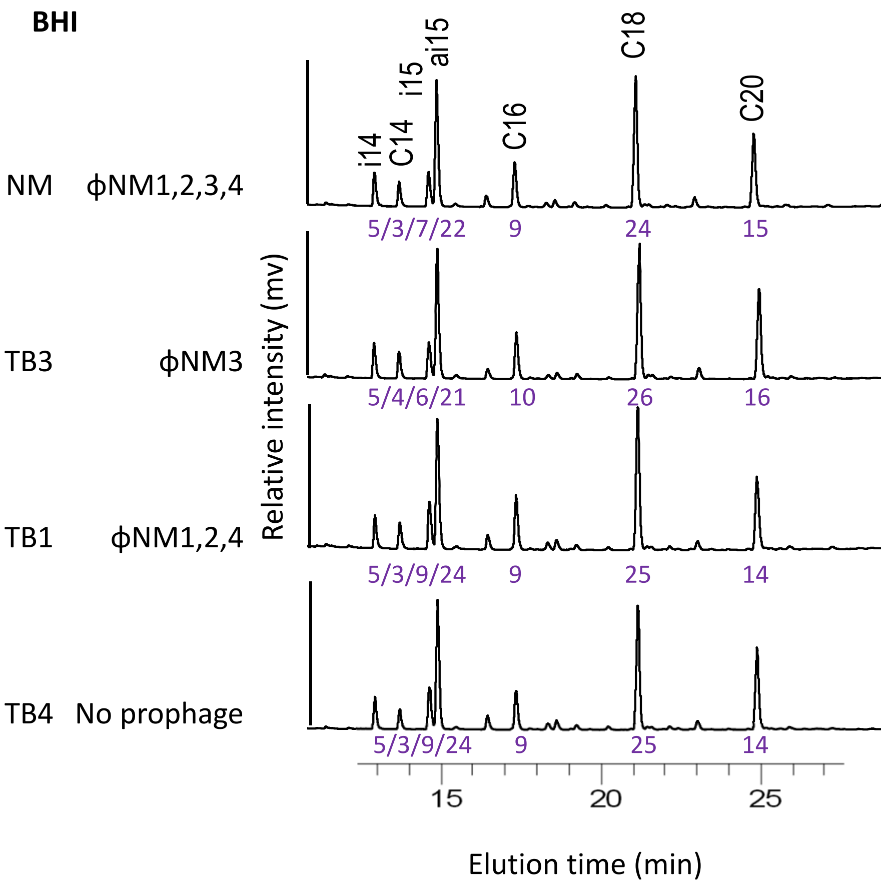


**Supplementary Figure S2.** FA profiles of *S. aureus* cultured in BHI medium are not affected by the prophage status. Four isogenic Newman strains, lysogenized or not by prophages as shown, were cultured in BHI. FAs were extracted and analyzed as described in Fig. 1B. Representative FA profiles and means of relative amounts (in purple) from three independent experiments are shown. Statistical significance was determined by the Kruskal-Wallis test to compare the FA compositions between the four strains. Profiles were not significantly different, p value> 0.05.

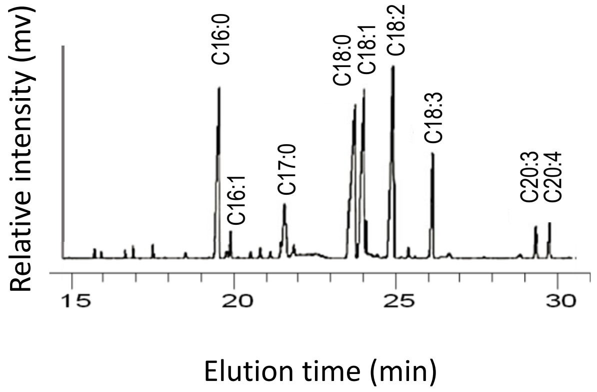


**Supplementary Figure S3.** FA profile from adult bovine serum. A representative profile determined as in Figure 1B is shown. This serum contains different PUFAs: linoleic acid (C18:2), α-linolenic acid (C18:3 ω-3), dihomo-γ-linolenic acid (20:3 ω−6), and arachidonic acid (C20:4 ω-6).

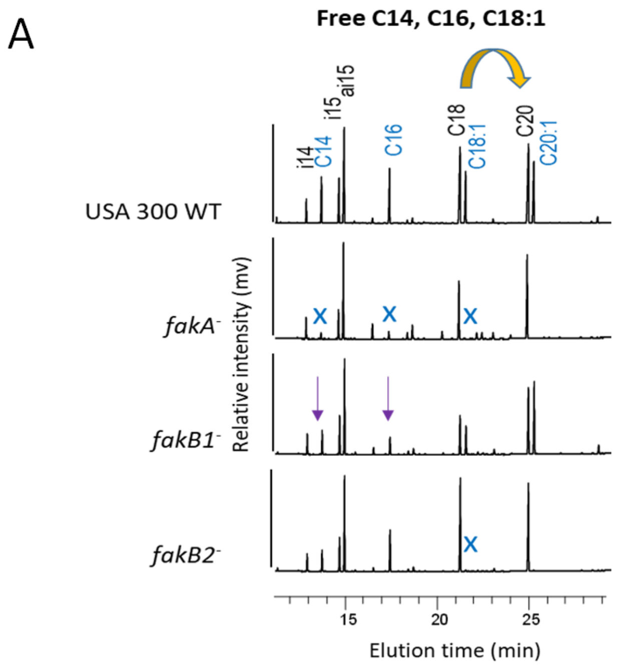


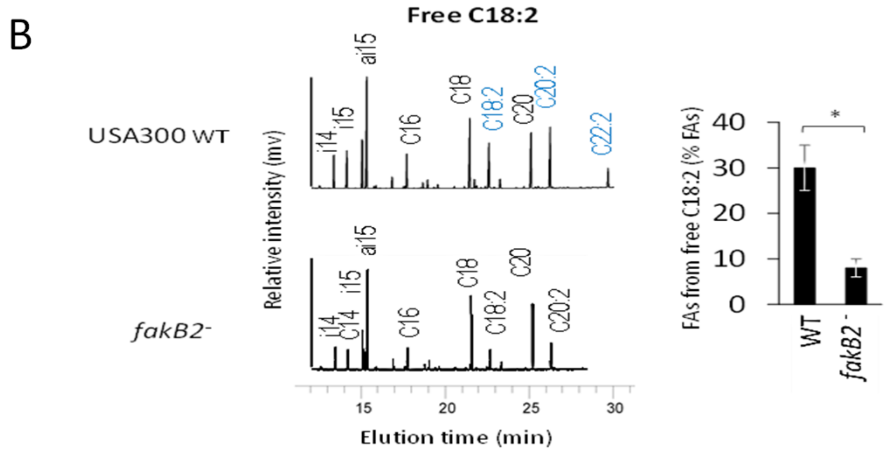


**Supplementary Figure S4.** Phenotype validation of *fakA, fakB1* and *fakB2* mutants. The USA300 wild type and mutant strains (Table 1) were cultured in BHI medium in the presence of exogenous FAs. Total membrane FAs were extracted, analysed and presented as in Fig. 1B. (A) FAs profiles from the WT strain and the *fak* mutant cultured in the presence of an FA cocktail (free C14:0, C16:0, and C18:1, 0.17 mM each). The three exogenous FAs and elongated forms are indicated in blue. Orange arrow indicates C18:1 elongation into C20:1. Blue crosses indicate that *fakA* is essential for FA incorporation and that *fakB2* is essential for C18:1 incorporation. Purple arrows indicate that *fakB1* is involved in incorporation of C14 and C16. (B) *fakB2* is required for incorporation of free C18:2. The WT strain and the *fakB2*::tn mutant were cultured in in the presence 10µM free C18:2. In profiles, C18:2 and its elongated forms are represented in blue. Note that free C18:2 is poorly incorporated in the *fakB2* mutant compared to the WT. Data presented are means +/- standard deviations from independent experiments (n=3). Statistical significance was determined by the Mann-Whitney test. *, p≤0.05.

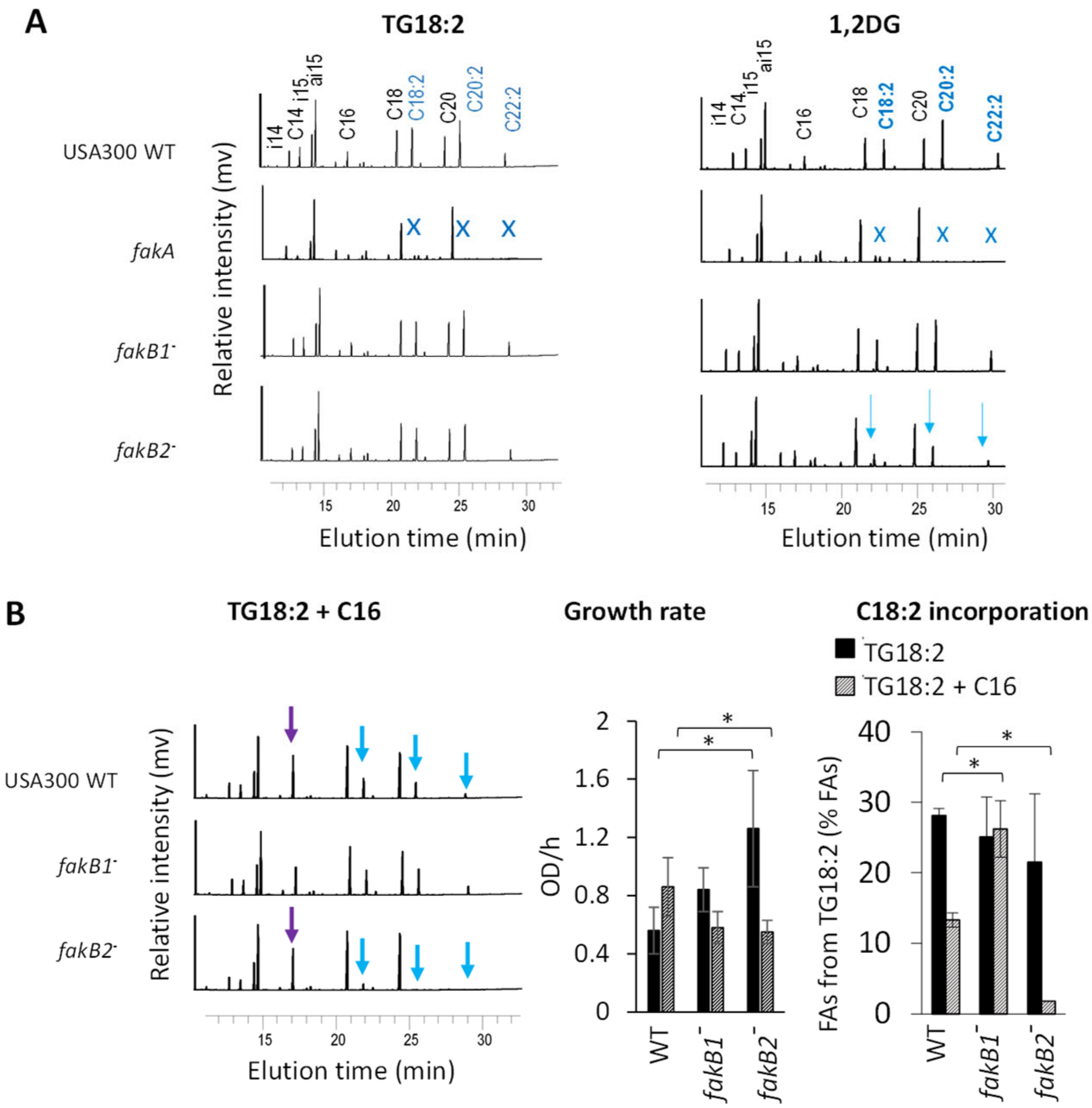


- **Supplementary Figure S5.** Roles of *fak* genes in the incorporation of C18:2 from TG18:2 or 1,2DG. Experiments and FAs profiles were performed as described in Figure 4, except that the USA300 WT and *fak* mutant strains (Table 1) were cultured in the presence of 30µM TG18:2 or 1,2DG, as sources of C18:2. (A) Representative FA profiles from *fakA, fakB1* and *fakB2* mutants cultured in the presence of TG18:2 or 1,2DG. Left: The *fakA* defective strain fails to incorporate C18:2 from TG18:2, while both *fakB1* and *fakB2* mediate incorporation of TG18:2-derived C18:2. Right: The *fakB2* gene, and not *fakB1*, affects C18:2 incorporation from 1,2 dilinolein (1,2DG). (B) C16 inhibits incorporation of C18:2 from TG18:2 *via* competition for FakB1. The *fakB2* mutant (encoding only *fakB1*) fails to incorporate TG18:2-derived C18:2 in the presence of C16. Growth rates presented as $OD_{600}$ per hour suggest that FakB1 protects from C18:2 toxicity and that C16 inhibits growth when only FakB1 is active by releasing free C18:2 in *S. aureus*. TG18:2-derived C18:2 and elongated forms are indicated in blue, and blue crosses indicate their absence. Purple arrows indicate a C16 increase due to exogenous supply. Blue arrows indicate inhibition of TG18:2-derived C18:2 and elongated forms. Profiles are representative of independent experiments (n=3). Histograms are means +/- standard deviations from these experiments. Statistical significance was determined by the Mann-Whitney test.; *, $p \leq 0.05$.

-
